# Improvements in the detection power of algorithms for analyzing next-generation sequencing based bulked segregant analysis data via estimating thresholds at the genomic region level

**DOI:** 10.1101/2023.03.12.532308

**Authors:** Jianbo Zhang, Dilip R Panthee

**Affiliations:** Mountain Horticultural Crops Research and Extension Center, Department of Horticultural Science, North Carolina State University, Mills River, NC 28759, USA

## Abstract

Next-generation sequencing based bulked segregant analysis (BSA-Seq) has been widely used in identifying genomic regions associated with a trait of interest. However, the most popular algorithms for BSA-Seq data analysis have relatively low detection power, and high sequencing depths are required for the detection of genomic regions linked to the trait. Here we estimated the confidence intervals/thresholds of the popular algorithms at the genomic region level and increased the detection power of these algorithms by at least 5 folds, which should drastically reduce the sequencing cost of BSA-Seq studies.

## Introduction

Bulked segregant analysis (BSA) was developed for rapidly identifying genetic markers linked to a trait of interest (Michelmore *et al*., 1991; Giovannoni *et al*., 1991). For a particular trait, a certain number of individuals of a segregating population are selected to make two bulks or pools that have contrasting phenotypes of the trait. Equal amounts of DNA are pooled from each individual within a group. The pooled DNA samples are then examined via different methods to identify bulk and parent-specific genetic markers. Collectively, the genomic region(s) controlling the trait should be different while the other unlinked regions should be the same or very similar between the bulks. Thus, the bulk and parent-specific markers should be associated with the trait. Initially, the bulks were genotyped using technologies such as restriction fragment length polymorphism (RFLP) and amplified fragment length polymorphism (AFLP) (Asnaghi *et al*., 2004; Cai *et al*., 2003; Hartl *et al*., 2011; Zhang *et al*., 2013; Michelmore *et al*., 1991; Giovannoni *et al*., 1991). This approach relies on the absence/presence of trait-associated DNA fragments in the gel, which works well for traits controlled by a single qualitative gene. Recently, next-generation sequencing has been used for genotyping the bulks (BSA-Seq) (Gu *et al*., 2017; Imerovski *et al*., 2019; Klein *et al*., 2018; Trick *et al*., 2012; Arikit *et al*., 2019; Cao *et al*., 2021; Q. Chen *et al*., 2018; Clevenger *et al*., 2018; Das *et al*., 2015; Daware *et al*., 2016; Takagi *et al*., 2013; Magwene *et al*., 2011; Ehrenreich *et al*., 2010). Genome-wide structural variants (SV) such as single nucleotide polymorphisms (SNP) and small insertions/deletions (InDel) can be easily identified via this approach, and SVs associated with different traits, including single gene-controlled traits, qualitative traits controlled by multiple genes with or without epistatic effects, and quantitative traits, can be easily detected with various algorithms.

An SV in a BSA-Seq dataset can be biallelic, triallelic, or tetraallelic. Biallelic SVs are the most abundant if the parents for generating the population are highly homozygous. Only this type of SV is used in most BSA-Seq analysis algorithms. For a biallelic SV locus, one of its alleles is termed the reference allele (REF), while the other is referred to as the alternative allele (ALT). When the parental genome sequences are available, the SV alleles from one of the parents are REF alleles, while the SV allele from the other parent are ALT alleles. Because each bulk contains many individuals, the vast majority of SV loci in a bulk contain both REF and ALT alleles. For each SV, the number of reads of REF/ALT alleles is termed allele depth (AD). Because of the phenotypic selection via bulking, for trait-associated SVs, their ALT allele should be enriched in one bulk while their REF allele should be enriched in the other. However, for SVs not associated with the trait, both of their ALT and REF alleles would be randomly segregated in both bulks and enriched in neither. Hence, these four AD values (REF/ALT reads from each bulk) can be used to assess how likely an SV is associated with the trait.

A few algorithms have been developed to identify trait-associated genomic regions based on BSA-Seq data. The allele frequency method (QTL-seq) is the most widely used for such a purpose (Takagi *et al*., 2013). It calculates the REF or ALT allele frequencies (AF) of an SV in both bulks with the four AD values and uses the allele frequency difference (ΔAF) between bulks to assess the likelihood the SV locus is associated with the trait; obviously, a high absolute value of the ΔAF suggests that the SV is more likely associated with the trait. The *G*-statistic method uses G-test to calculate the *G*-statistic value of an SV with the four AD values and uses the calculated value to judge how likely the SV is associated with the trait, a high *G*-statistic value suggests that the SV is more likely associated with the trait (Magwene *et al*., 2011). We have previously developed the significant structural variant method for BSA-Seq data analysis (Zhang and Panthee, 2020, 2022). In this method, an SV is assessed with Fisher’s exact test using its AD values of both bulks. An SV with a low p-value of Fisher’s exact test tends to have a high absolute ΔAF and *G*-statistic values and is more likely linked to the trait. An SV is considered significant if the p-value of Fisher’s exact test is lower than a specific type I error rate, e.g., α=0.01, and non-significant otherwise. Since SVs normally are not evenly distributed across chromosomes, we use the ratio of the significant SVs (sSV) to the total SVs (totalSV) within a genomic region to judge if this region is associated with the trait of interest in our method. A genomic region with a high sSV/totalSV ratio is more likely associated with the trait. The sSV/totalSV ratio is a parameter at the genomic region level, whereas both the allele frequency and the *G*-statistic value are parameters at the SV level, which is the key difference between the significant structural variant method and the standard allele frequency and *G*-statistic methods, and the root cause of our method’s higher statistical power in the detection of the genomic region-trait associations.

The *G*-statistic and allele frequency methods (these are referred to as the standard methods hereafter) were developed in 2011 and 2013, respectively, much earlier than the significant structural variant method. As such, these methods became the established norm for BSA-Seq analysis in many laboratories. Although our method has higher detection power, the standard methods are still widely used in recent publications. Due to the prevalence of their use, it is necessary to improve the detection power of the standard methods if possible. In this study, we estimated the threshold of the standard methods at the genomic region level, which increases the detection power of these methods to a level similar to that of the significant structural variant method.

## Materials and Methods

The rice sequencing data used in this study were generated by (Lahari *et al*., 2019). In that study, the size of the F_2_ population was 178, and the size of both the resistant and the susceptible bulk were 23 plants each. The parents of this population are LD24 and VialoneNano. Illumina MiSeq Sequencing System and MiSeq v3 chemistry were used for sequencing the DNA samples of both the parents and the bulks. The accession numbers of the sequences of LD24, VialoneNano, the resistant bulk, and the susceptible bulk are ERR2696318, ERR2696319, ERR269632, and ERR2696322, respectively.

Most methods used in this study had been described previously (Zhang and Panthee, 2020, 2022), minor modifications are described as below.

### SV calling

The rice BSA-Seq sequencing data were downloaded from the European Nucleotide Archive (ENA) using the Linux program wget, and the rice reference sequence (Release 47) was downloaded from https://plants.ensembl.org/Oryza_sativa/Info/Index. Sequencing data preprocessing and structural variant calling were performed using fastp, samtools, BWA-MEM, and GATK4 (Li *et al*., 2009; Li, 2011, 2013; McKenna *et al*., 2010; S. Chen *et al*., 2018) as described previously (Zhang and Panthee, 2020, 2022). The SV calling-generated .vcf files were converted to .tsv files using the GATK4 utility VariantsToTable, which served as input files for our Python script.

### Workflow of the Python scripts

The SV dataset generated via SV calling was processed with our Python script to identify significant SV-trait associations. A single script contains all three algorithms and the code is available on http://github.com/dblhlc. The workflow of the scripts is as follows:

1. Convert the .tsv/.csv input file(s) generated via SV calling to a pandas DataFrame (size-mutable and potentially heterogeneous two-dimensional tabular data structure with labeled rows and columns) (McKinney 2010; Reback *et al*. 2021).
2. Filter SVs on the pandas DataFrame. Details of SV filtering can be found in our previous publication (Zhang and Panthee, 2022).
3. Perform Fisher’s exact test to obtain the p-value, calculate the ΔAF value, or perform G-test to obtain the *G*-statistic value of each SV in the filtered pandas DataFrame using its four AD values (AD_ref1_ and AD_alt1_ of bulk 1 and AD_ref2_ and AD_alt2_ of bulk 2). The p-values of Fisher’s exact tests, the ΔAF values, and the *G*-statistic values of all SVs on each chromosome are smoothed by applying a Savitzky-Golay filter (Savitzky and Golay, 1964).
4. Estimate the ΔAF and the *G*-statistic thresholds of each SV in the filtered pandas DataFrame via simulation. A Savitzky-Golay filter (Savitzky and Golay, 1964) is applied to these values at the chromosome level as well to smooth the threshold curves. The thresholds will be used to identify the significant peaks/valleys in the plots generated in the next step in the old approach.
5. Estimate genome-wide thresholds for all three BSA-Seq algorithms.
6. Use the sliding window algorithm to plot the sSV/totalSV ratios, the ΔAF values, or the *G*- statistic values against their genomic positions.
7. Identify potential significant peaks/valleys using the genome-wide thresholds.
8. Calculate the thresholds of sliding windows with a potential significant peak/valley to verify if the potential peaks/valleys are really significant.

### Sliding window settings

The sliding windows algorithm was used to mitigate noise generated by random errors introduced in bulking and sequencing. For all three methods, the size of the sliding windows is 2 Mb and the incremental step is 10 kb.

### Simulate allele depth values under the null hypothesis

If a biallelic SV is not associated with the trait, the probability of its ALT allele in the bulks is 0.5 for an F_2_ population and 0.25/0.75 for a backcross population, and its AD_alt_ should follow the binomial distribution. Based on that, we used the above probability and the sequencing depth (AD_ref_+AD_alt_, number of trials) of an SV to simulate the allele depths of its REF/ALT alleles in a bulk. The simulated smAD_ref1_/smAD_alt1_ of bulk 1 and smAD_REF2_/smAD_ALT2_ of bulk 2 are used for threshold estimation below.

### Calculation of ΔAF/*G*-statistic and estimation of their thresholds

Calculation of the *G*-statistic value and the ΔAF value of each SV in the dataset and estimation of its thresholds were performed as described previously (Zhang and Panthee, 2020, 2022). In brief, equation 1 is used for ΔAF calculation while equation 2 is used for *G*-statistic calculation. In equation 2, *O* indicates the AD value of an SV (AD_REF1_, AD_ALT1_, AD_REF2_, or AD_ALT2_), *E* is the expected AD value under the null hypothesis (the SV is not linked to the trait) and is calculated as in the original *G*-statistic method (Magwene *et al*., 2011), and *ln* denotes the natural logarithm. Most sliding windows contain many SVs. The ΔAF/*G*-statistic value of a sliding window is the mean of the ΔAF/*G*-statistic values of all SVs in it.

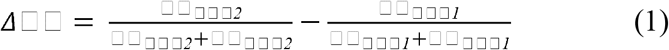

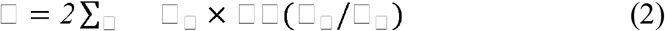

ΔAF/*G*-statistic threshold estimation for each SV in the dataset is performed by simulation. For each SV, its ΔAF or *G*-statistic value is calculated using the simulated AD values in both bulks with equation 1 or equation 2. This process is repeated 10,000 times, the threshold for the allele frequency method is the 99% confidence interval of the 10,000 ΔAF values, and the threshold for the *G*-statistic method is the 99.5^th^ percentile of the 10,000 *G*-statistic values. As in the real dataset, the threshold of a sliding window is the average value of all SVs in it.

### Identification of significant SVs

For each SV in the dataset, Fisher’s exact test was performed using its four AD values. An SV with its p-value less than ⟨=0.01 is defined as a significant SV. Calculating the sSV/totalSV ratio of a genomic region is straightforward.

### Genome-wide and sliding window specific threshold estimation

Calculating a threshold for all sliding windows of the SV dataset would take a very long time. To overcome this obstacle, we first calculate a genome-wide threshold and use it to identify potential significant peaks/valleys, then sliding window thresholds of these peaks are calculated via simulation to verify if these peaks/valleys sliding windows are really significant.

#### Genome-wide threshold

The number of SVs that are the same as the average number of SVs per sliding window are randomly selected from the entire dataset. For each SV in this sample, the four simulated AD values are used to obtain the p-value via Fisher’s exact test and to calculate the ΔAF/*G*-statistic values. An SV with its p-value less than ⟨=0.05 is considered an sSV (⟨=0.01 is used for the real dataset to reduce the chance of false positives). Then the ΔAF, *G*-statistic, and sSV/totalSV ratio of the sample (treated as a genomic region) are calculated as above. This process (starting from sampling SVs) is repeated 10,000 times. The 99% confidence interval of the simulated 10,000 ΔAF values is used as a significant threshold for the allele frequency method, and the 99.5^th^ percentile of these 10,000 simulated *G*-statistic values or sSV/totalSV ratios is used as the significance threshold for the significant structural variant or *G*-statistic methods for detection of potential significant peaks.

#### Sliding window threshold

Estimating a sliding window specific threshold is very similar to estimating the genome-wide threshold; the only difference is that we do not need to sample SVs from the genome. Instead, the SVs and their associated AD values in each bulk in the sliding window are used for simulation, and the rest is the same as above.

## Results

### Appropriate threshold estimation of BSA-Seq data analysis

If an SV or a genomic region is not associated with a trait, its ΔAF, *G*-statistic, and sSV/totalSV values should be close to zero (base value). Otherwise, due to allele enrichment/depletion, the values of these variables for an SV or a genomic region will deviate from the base value significantly, and thus peaks/valleys will be formed in the trait-associated genomic regions in the BSA-Seq curve. However, chance variation (random error) can be introduced during bulking and sequencing, which can lead to sequencing depth variations across the genome and result in the formation of false peaks/valleys. The genuine peaks/valleys should have higher absolute values than those of the false peaks/valleys. An appropriate confidence interval/threshold is needed to distinguish these two types of peaks/valleys.

Confidence intervals/thresholds should be estimated at an appropriate level. If we try to associate an SV with a trait, the confidence intervals/thresholds should be estimated at the SV level. In contrast, if we try to associate a genomic region with a trait, the thresholds should be estimated at the genomic region level. We can associate an SV to a trait with an extremely large population, large bulk sizes, and high sequencing coverage. Large populations ensure the occurrence of recombination events between the trait-controlling gene and its flanking sequences, a large bulk size ensures the inclusion of such recombination events in the bulks for genotyping, and high sequencing coverage allows us to differentiate the trait-controlling SVs from the flanking ones and identify the trait-associated SV with sufficient statistical significance. However, doing so is probably not feasible because of the labor requirement and high costs. Therefore, it is more realistic to associate a genomic region(s) to a trait of interest. In this case, instead of estimating the confidence interval/threshold at the SV level, it is more appropriate to estimate it at the genomic region level.

Confidence intervals/thresholds are estimated in different ways in different algorithms. The most widely used allele frequency method calculates the confidence interval of each SV in a dataset via simulation and uses the upper/lower bound means of the confidence intervals of all SVs in a genomic region to judge if this region is significant (Takagi *et al*., 2013). The original *G*-statistic method uses a false discovery rate approach (Benjamini and Hochberg, 1995; Benjamini and Yekutieli, 2001; Davies and Gather, 1993) to estimate the genome-wide threshold and the threshold was relatively low when a yeast dataset was analyzed (Magwene *et al*., 2011). But the threshold was much higher when a rice dataset (Yang *et al*., 2013) was analyzed, and thus its detection power was similar to that of the allele frequency method (Mansfeld and Grumet, 2018). We used simulation to estimate the *G*-statistic threshold of each SV in the same rice dataset in a way similar to that in the original allele frequency method, and the results revealed that the threshold obtained via our simulation approach is less conservative than that obtained via the false discovery rate approach (Mansfeld and Grumet, 2018). The sSV/totalSV ratio used in our significant structural variant method is a parameter at the genomic region level, its threshold is estimated at the genomic region level via simulation, and our method has at least five times higher detection power than the standard methods (Zhang and Panthee, 2020).

We simulate the ALT/REF allele depths of each SV based on the null hypothesis when estimating confidence interval/threshold. Under the null hypothesis, the ALT/REF allele frequencies of most SVs in a genomic region should be centered around the probability of the ALT/REF alleles, and only a small potion of SVs have extreme allele frequencies; moreover, the distributions of the allele frequencies should be the same or very similar between the two bulks. However, the confidence interval/threshold of an SV are extreme ΔAF/*G*-statistic values; it is not appropriate if we calculate the ΔAF confidence interval and *G*-statistic threshold of each SV in a genomic region and use their mean as the confidence interval/threshold of this genomic region.

Doing so would assume that the allele frequencies of all SVs in the genomic region are at the extreme ends of the allele frequency distribution and very different from the probability of the REF/ALT alleles, which is against the null hypothesis, and thus can lead to very conservative confidence interval/threshold values.

### The confidence interval/threshold at the genomic region level is very different from that at the SV level

We analyzed a randomly selected sliding window from the rice nematode resistance dataset (Lahari *et al*., 2019) to test how confidence interval/threshold estimation behaves at the SV and genomic region levels. Since slightly better detection power was achieved in estimating the *G*-statistic thresholds via simulation than via the false discovery rate approach with the same dataset (Zhang and Panthee, 2020; Mansfeld and Grumet, 2018), we used simulation to estimate both the ΔAF confidence intervals and the *G*-statistic threshold here. The selected sliding window is on chromosome 1 and consists of 7343 SVs in this case. We simulated the allele depth value of each SV in the sliding window once under the null hypothesis and calculated its corresponding ΔAF and *G*-statistic values (Figure 1A and 1 B). As the way in which the real sliding window ΔAF value and *G*-statistic value are calculated, the simulated ΔAF value and *G*-statistic value of the sliding window are the mean of these simulated ΔAF values and the *G*-statistic values of SVs, respectively. As expected, the simulated sliding window ΔAF value was very close to zero and the majority of the simulated SV ΔAF values were centered around zero (Figure 1A), because the two bulks should be the same under the null hypothesis. Then we estimated the confidence interval and threshold of this sliding window at the SV level as described previously (Zhang and Panthee, 2020, 2022). The ΔAF confidence interval and the *G*-statistic threshold of each SV in the sliding window were obtained by simulation under the null hypothesis, and the distribution of these confidence intervals and thresholds were presented in

**Figure 1.**
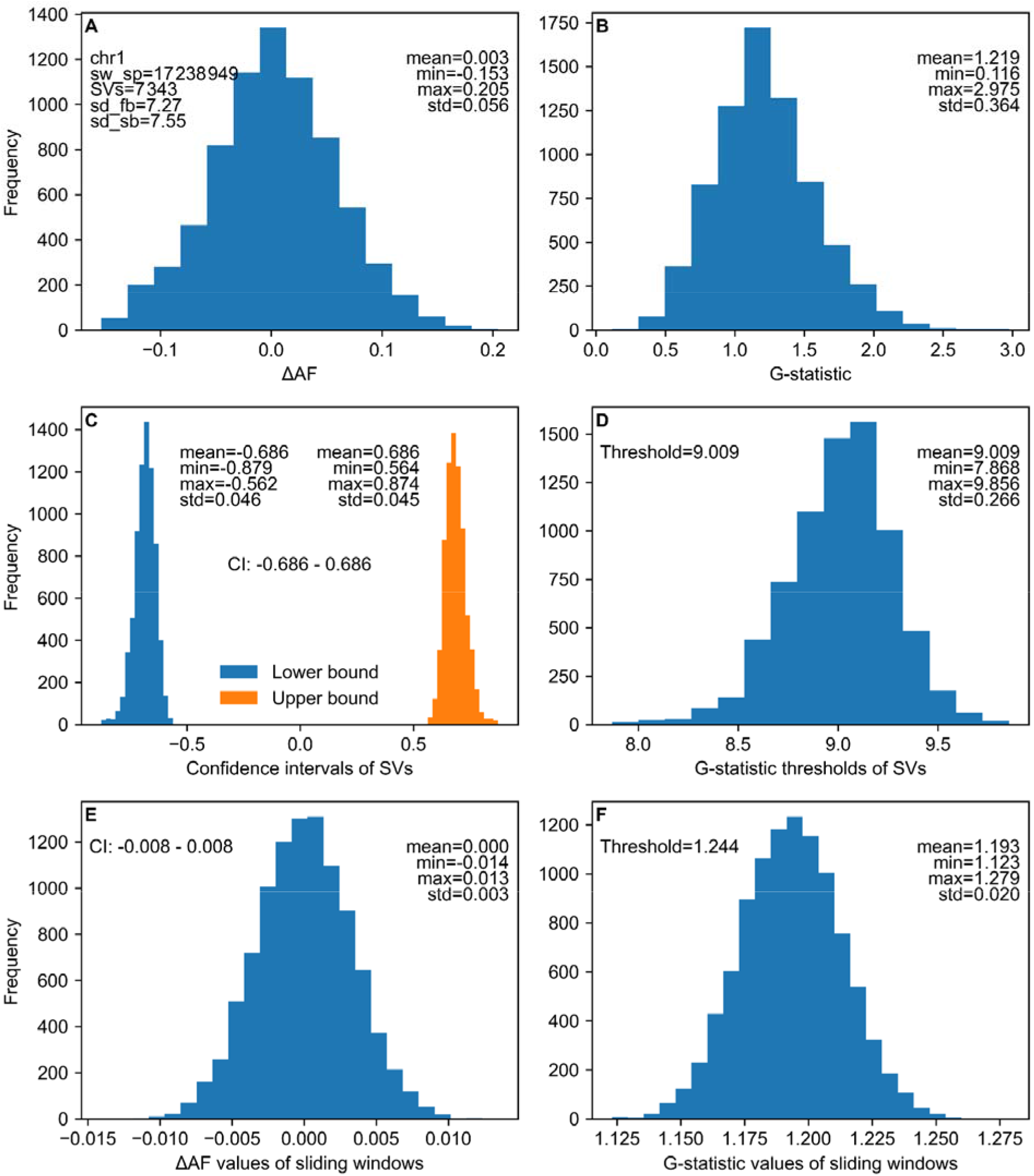
Distributions of ΔAF and *G*-statistic related values in a randomly selected sliding window. The sliding window is on chromosome 1; it consists of 7343 SVs, the average sequencing depth in the first bulk is 7.27 while the average sequencing depth is 7.55 in the second bulk. **A**. Histograms of the simulated ΔAF values of SVs in the sliding window. **B**. Histograms of the simulated *G*-statistic values of SVs in the values in the sliding window. **C**. Histograms of the ΔAF confidence intervals of SVs in the sliding window. **D**. Histograms of the *G*-statistic thresholds of SVs in the sliding window. **E**. Histogram of 10000 simulated ΔAF values of the sliding window. **F**. Histogram of 10000 simulated *G*-statistic values of the sliding window.. All simulations were performed under the null hypothesis. The meanings of the abbreviations used here: sw_sp - the sliding window start point; SVs - the number of SVs in the sliding window; sd_fb - the average sequencing depth of the SVs in the sliding window in bulk1; sd_sb - the average sequencing depth of the SVs in the sliding window in bulk2.

Figures 1C and 1D, respectively. The upper bound of the ΔAF confidence interval of the sliding window is the mean of the upper bounds of all SVs and the lower bound of the ΔAF confidence interval of the sliding window is the mean of the lower bounds of all SVs in the sliding window. Likewise, the *G*-statistic threshold of the sliding window is the mean of the *G*-statistic thresholds of all SVs in the sliding window.

To estimate the confidence interval/threshold of this sliding window at the sliding window level, we repeat the process in Figures 1A and 1B for 10,000 times to obtain 10,000 corresponding sliding window ΔAF value and *G*-statistic values, then we use ⟨=0.01 (two-tailed) to find the ΔAF confidence interval of the sliding window and use ⟨=0.01 (one-tailed) to find the *G*-statistic threshold of the sliding window (Figures 1E and 1F). The results revealed that the ΔAF confidence interval is much drastically narrower at the sliding window level (−0.008 - 0.008 vs - 0.686 - 0.686), and the *G*-statistic threshold is much lower at the sliding window level as well (1.244 vs 9.009). As long as a sliding window does not contain too few SVs, most of its simulated ΔAF values will be very close to zero. Thus it is likely that the thresholds of the sliding window will be too far away from zero, which also suggests that the background noise of the allele frequency method can be very low.

### Estimation of confidence intervals/thresholds at the genomic region level greatly increase the detection power of the standard methods

Using the approaches described above, we reanalyzed the rice nematode resistance data (Lahari *et al*., 2019) to test how threshold estimation at the sliding window level affects the detection power of the standard algorithms. We first used a genome-wide confidence interval/threshold to identify potentially significant peaks/valleys, then used sliding windows specific confidence intervals/thresholds to test if these peaks/valleys were really significant. Previously we used α=0.01 to identify sSVs in the real dataset and used α=0.1 to identify sSVs in the simulated dataset. In this way we obtained higher thresholds and lowered the chance of false positives.

Here we used α=0.01 to identify sSVs in both real and simulated dataset to level the play field for the detection power comparison among different algorithms. Only the major peak on chromosome 11 was identified as significant using the old standard methods (Lahari *et al*., 2019; Zhang and Panthee, 2022). However, in addition to this major peak, many minor peaks on chromosomes 1, 3, 4, 5, 6, 7, 8, 9, 10, 11, and 12 become significant as well when the thresholds are estimated at sliding window level (Figure 2), which means that the detection power of the standard methods is increased by at least 5 folds (Zhang and Panthee, 2020, 2022), because we used less stringent threshold for the significant structural variant method here. The average sliding window threshold of the *G*-statistic is 9.00585070076413, and the average sliding window confidence interval of the allele frequency method is -0.68221874129103-0.683282282173592 using the old approach. Whereas these values are 1.2492238068569297 and -0.00979694695809328 to 0.009111848721497237, respectively, using the new approach, very close to their base values. The genome-wide threshold of the significant structural method is 0.004921092821992194. Using the new approach, the threshold value is drastically lower and the confidence interval is drastically narrower, thus the detection power of the standard methods is dramatically increased.

**Figure 2.**
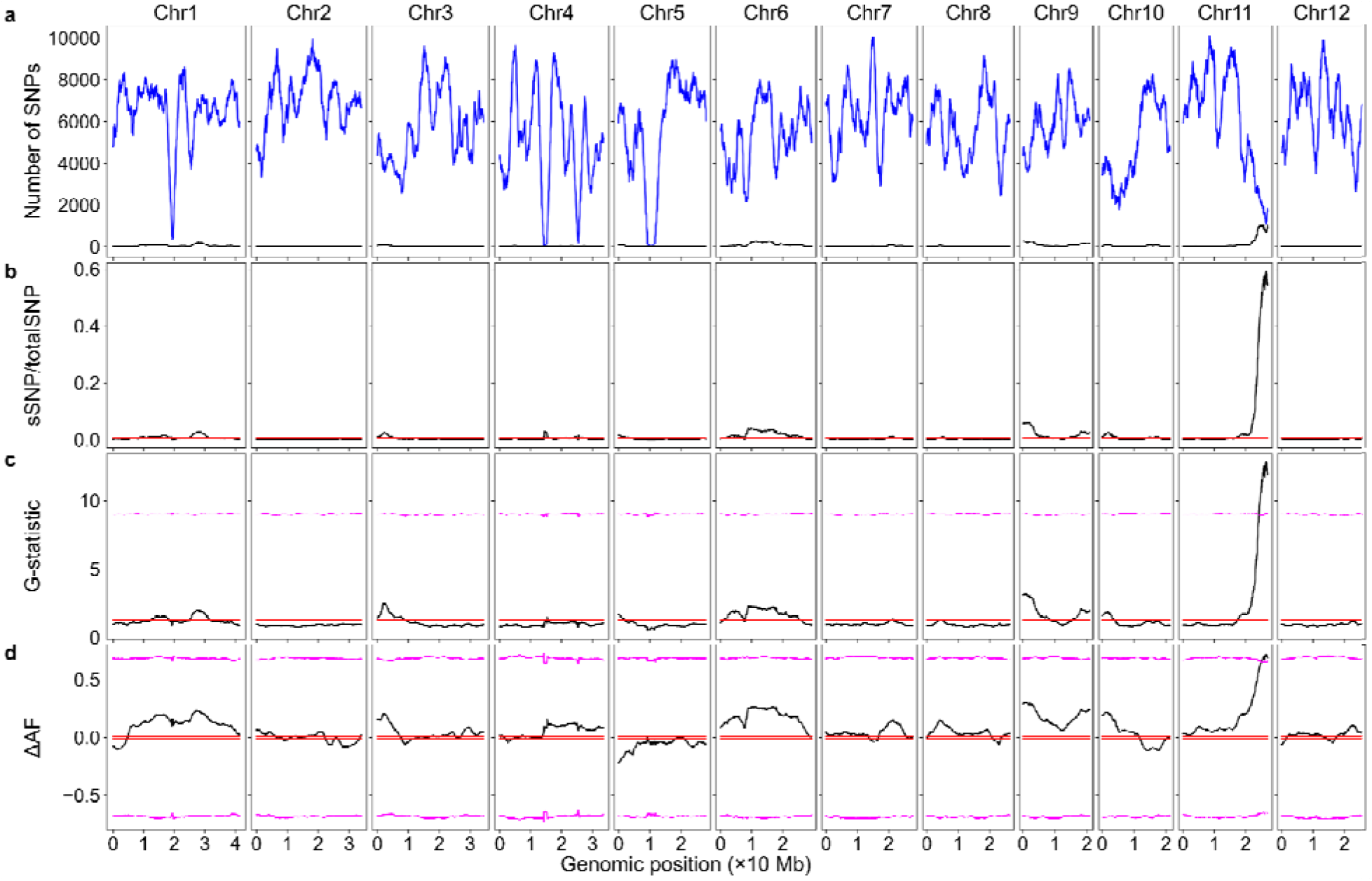
BSA-Seq data analysis using both the parental and bulk genome sequences in rice. The red lines/curves are genome-wide thresholds, while the magenta lines/curves are the thresholds estimated at the SV level. The black curves represent the number of sSVs (A), sV/totalSV ratios (B), *G*-statistic values (C), or ΔAF values (D). (A) Genomic distributions of sSVs and total SVs. Blue curves: total SVs. (B) Genomic distributions of sSV/totalSV ratios. (C) Genomic distributions of *G*-statistic values. (D) Genomic distributions of ΔAF values.

When estimating thresholds at the genomic region level, the *G*-statistic method has lower detection power while the significant structural variant and allele frequency methods have similarly higher detection powers. However, all the additional significant peaks/valleys detected via the allele frequency or the significant structural variant methods are extremely minor.

Although the way we obtain the new thresholds is statistically sound, it is very likely the very minor peaks/valleys do not contribute much to the trait phenotype; moreover, lower thresholds may increase the chance of false positives. It is possible that some very minor genuine peaks/valleys might be missed, slightly increasing the threshold can decrease the chance of false positives, which should be practically beneficial. We used α=0.01 for the identification of the significant structural variants in both the real and simulated data here. If we use α=0.05 as described in the Methods section or α=0.1 as we did previously for the identification of the significant SVs in simulated data, and use α=0.01 for the identification of the significant SVs in real data, the threshold of the significant structural variant method could be significantly higher. The ΔAF value of an SV will fluctuate closely around zero under null hypothesis (it is not associated with the trait); in a genomic region, positive values and negative values would cancel each other when calculating the ΔAF mean, leading to the threshold value very close to zero.

Thus, we will use the absolute ΔAF of each SV in a genomic region to calculate the ΔAF of this region. If the cutoff ΔAF of these 10000 values at α=0.005 (one-tailed) is *a*, the confidence interval will be between -*a* and *a*. These approaches will be used to estimate the thresholds of the allele frequency and significant structural variant (α=0.05) methods for sections below.

The sliding window with the major peak on chromosome 11 contains 1139 SVs; the average sequence depths of these SVs are 7.6540 and 8.7471, respectively, in bulks 1 and 2, and the corresponding standard deviations are 3.0440 and 2.8205. The threshold of its *G*-statistic was decreased from 9.0472 to 1.2930, and the confidence interval of its allele frequency difference (ΔAF) method was narrowed down from -0.6500-0.6496 to -0.2229-0.2229. To test the precision of the new threshold estimation approach, we repeated the process 500 times and generated 500 threshold values of the sliding window containing the major peak on chromosome 11 for all three methods. These threshold/confidence interval values are extremely consistent, and the coefficient of variance (CV) values of the 500 values are less than 0.01 for the standard methods and 1.16% for the significant structural variant method. But the range of the 500 thresholds of the latter is very narrow, their min is 0.0360 and their max is 0.0369 (Table 1).

**Table 1.** Statistical summary of the 500 thresholds of different algorithms.

| | sSV/totalSV | $G$-statistic | $\Delta AF$ |
| --- | --- | --- | --- |
| Min | 0.0359964881474978 | 1.2858852296992 | 0.22229617283634 |
| Max | 0.0368744512730465 | 1.30044138260958 | 0.223570624666076 |
| Mean | 0.036314503950834 | 1.29299772054259 | 0.222924623486806 |
| SD | 0.000422208343043444 | 0.00247340791486133 | 0.000239143546550072 |
| CV | 0.011626438395388 | 0.00191292519357527 | 0.00107275518877001 |

### Identify significant peaks/valleys using *t*-test

SVs in genomic regions that are not linked to the trait of interest segregate randomly in both bulks, and thus the allele frequencies of the SVs in these regions should be similar between the bulks. However, due to phenotypic selection, the trait-associated genomic regions are enriched in one bulk, and depleted in the other bulk if the trait is not controlled by many epistatic genes, so the allele frequencies of the SVs in these regions should be significantly different between the bulks. A sliding window generally contains hundreds or thousands of SVs. If the mean of allele frequencies of a sliding window are significantly different between the bulks, it is very likely that this sliding window is associated with the trait. Thus *t*-test is an appropriate tool for the identification of such genomic regions. We performed paired *t*-tests (since the bulks share the same set of SVs) to verify if the potentially significant peaks/valleys identified above are really significant, and the results are very similar to those obtained via the allele frequency method and the significant structural variant method (supplementary file 1).

## Discussions

Datasets that do not contain trait-associated SVs are used for confidence interval/threshold estimation in both the allele frequency and *G*-statistic method. In the original *G*-statistic method, Magwene and colleagues (2011) used the natural logarithm smoothed *G*-statistic value (*ln(G’)*) of each SV in the dataset, the median absolute deviation of each *ln(G’)*, and an empirical constant to filter out outliers (trait-associated SVs) in the real dataset. This approach worked fairly well when analyzing a yeast dataset, and the obtained genome-wide threshold was relatively low. But no outliers could be filtered out using the same approach and the same empirical constant when a rice dataset was analyzed, and the threshold was high even after outliers were filtered out using a different approach (Mansfeld and Grumet, 2018). In the allele frequency method, the datasets without trait-associated SVs are generated via simulation. The probabilities of the ALT/REF alleles under the null hypothesis are known for a genetic segregating population, they are 0.5/0.5 for an F_2_ population and 0.25/0.75 for a backcrossed population; the sequencing depths of each SV in both bulks can be found in the real dataset. Simulation of the ALT/REF allele depth of each SV is just like a coin flip simulation that has been tested thoroughly by the statistics community. Therefore, we used the simulation approach for confidence interval/threshold estimation in this study.

Although very similar results were obtained via estimating confidence intervals/thresholds at the genomic region level or via *t*-test, we prefer the *t*-test approach. Performing *t*-tests on allele frequency mean difference between the bulks is logically simple and takes much shorter time than simulation. In addition, the *t*-statistic value and the *p* value obtained via *t*-test are always the same for the same sliding window, no matter how many times the test is performed. In contrast, confidence intervals/thresholds obtained via simulation vary from time to time, although the changes are very minor (Table 1); peaks that are very close to confidence intervals/thresholds can be identified as either significant or not significant if the script is run multiple times.

However, the *t*-test approach will not work if the parental genome sequences are not available, and we can use only the simulation approach for confidence interval/threshold estimation in this case.

SVs are not distributed evenly across a genome and the number of SVs in sliding windows (sliding window size) vary greatly from sliding window to sliding window. The sequencing depths are different in different sliding windows as well (Zhang and Panthee, 2020, 2022). A genome-wide confidence interval/threshold would be appropriate and more efficient for the identification of trait-associated genomic regions if neither the sliding window size nor the sequencing depth affects the estimation of the confidence interval/threshold. Otherwise, a sliding window specific confidence interval/threshold is preferred for the reliable identification of trait-associated genomic regions. The results presented in the supplementary file 1 showed that each sliding window has a different confidence interval/threshold value. To figure out how sliding window size and sequencing depth affect confidence interval/threshold estimation, we estimated confidence intervals/thresholds with different sliding window sizes or at different sequencing depth levels. Our results revealed that a large sliding window narrows/decreases the confidence interval/threshold of all three algorithms, while high sequencing depth narrows/decreases the confidence interval/threshold of the standard methods, but increases the threshold of the significant structural variant method (supplementary file 2). However, this does not mean that estimation of threshold at the genomic region level decreases the detection power of the significant structural variant method. The number of sSVs increases more in trait-associated genomic regions in real datasets with higher sequencing depths because of the way p-values are calculated in Fisher’s exact test, and thus increasing sequencing depth increases the detection power of the significant structural variant method as well (Zhang and Panthee, 2020, 2022).

Moreover, high sequencing depths result in the allele frequencies of SVs closer to their probability and the allele frequency mean of a sliding window with a large number of SVs tends to be more precise. Thus both sequencing depth and sliding window size can affect the *t*-test results and the significance status of a sliding window as well in the *t*-test approach. Therefore, a *t*-test or a sliding window specific confidence interval/threshold is preferred to judge if a peak/valley is significant. The whole-genome threshold approach used in the original *G* statistic method (Magwene *et al*., 2011) and the block regression mapping method (Huang *et al*., 2020) should be avoided unless the *t*-test requirements cannot be satisfied or it is not possible to obtain a sliding window specific confidence interval/threshold.

## Acknowledgements

We thank Irene E. Palmer for editing the manuscript.

