## supplementary file 2 for "Improvements in the detection power of algorithms for analyzing next-generation sequencing based bulked segregant analysis data via estimating thresholds at the genomic region level"

### **Sliding window sizes are negatively correlated with thresholds**

The results in the supplementary file 1 indicate each sliding window has a different confidence interval/threshold. Here we used simulation to test how the sliding window size and the sequencing depth affect the estimation of confidence interval/threshold. We first tested how sliding window sizes affect threshold estimation at the sliding window level. Sliding window sizes of 1000, 2000, 3000, and 4000 were tested with average sequencing depth of 15. We simulated sequencing depths with cv=0.4 (based on the rice nematode resistance date). The sequencing depth and the REF/ALT reads of each SV are simulated as described in the Methods section. Figure S1 shows the distributions of 10,000 values of sSV/totalSV, *G*-statistic, and allele frequency of a sliding window. With a fixed average sequencing depth (15), a larger window size resulted in smaller threshold values for all the methods. The means of these 10.000 values are very similar between different window sizes, however, the standard deviation becomes smaller with a larger sliding window size. Thus, a larger sliding window has less threshold value when the average sequencing depth is fixed.

### **Sequencing depths are reversely correlated with thresholds for the standard methods, but positively correlated with thresholds for the significant variant methods**

Next, we tested how sequencing depths affect threshold estimation. Sequencing depths of 5, 10, 15, and 20 were tested. Again sequencing depths were simulated with CV=0.4. We used the sliding window size of 6000 to test the effects of sequencing depth on threshold estimation. As shown in Figure S2, increasing sequencing depth slightly increases the threshold of the significant structural variant method but decreases the thresholds of the other methods. These results suggest that increasing sequencing depth can increase the detection power of the standard methods; however, it does not mean that doing so will decrease the detection power of the significant structural variant method. The number of sSVs increases more with higher sequencing depths in trait-associated genomic regions in real datasets because of the way p-values are calculated in Fisher’s exact test, and thus increasing sequencing depth increases the detection power of the significant structural variant method as well (Zhang and Panthee, 2020, 2022).

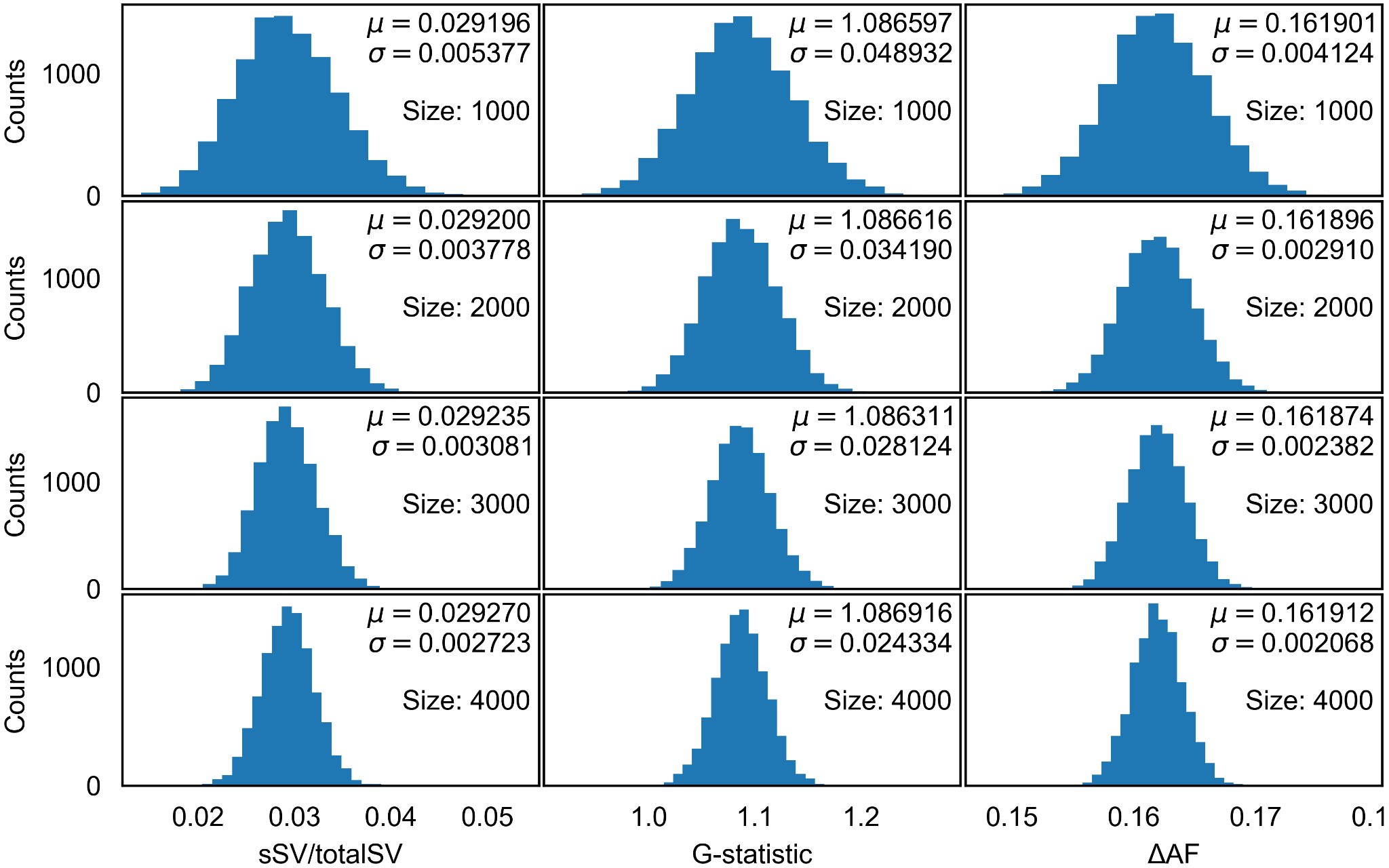

**Figure S1. Distribution of 10000 values of sSV/totalSV, *G*-statistic, and ΔAF obtained via simulation under the null hypothesis** (all SVs are not associated with the trait). Conditions: sequencing depth = 15; the coefficient of variation (sequencing depth) is 0.4, the number of SVs per sliding window - 1000, 2000, 3000, and 4000 (from top to bottom).

We generated 500 thresholds for each treatment in Figures 32 and 43. As for the nematode dataset, these threshold values are extremely consistent, and the CV values for all the treatments are less than 0.01. Analysis of variation (ANOVA) and Tukey’s Honest Significant Difference (HSD) tests revealed that the threshold difference between different sequencing depth/sliding window size levels is significant at α=0.01.

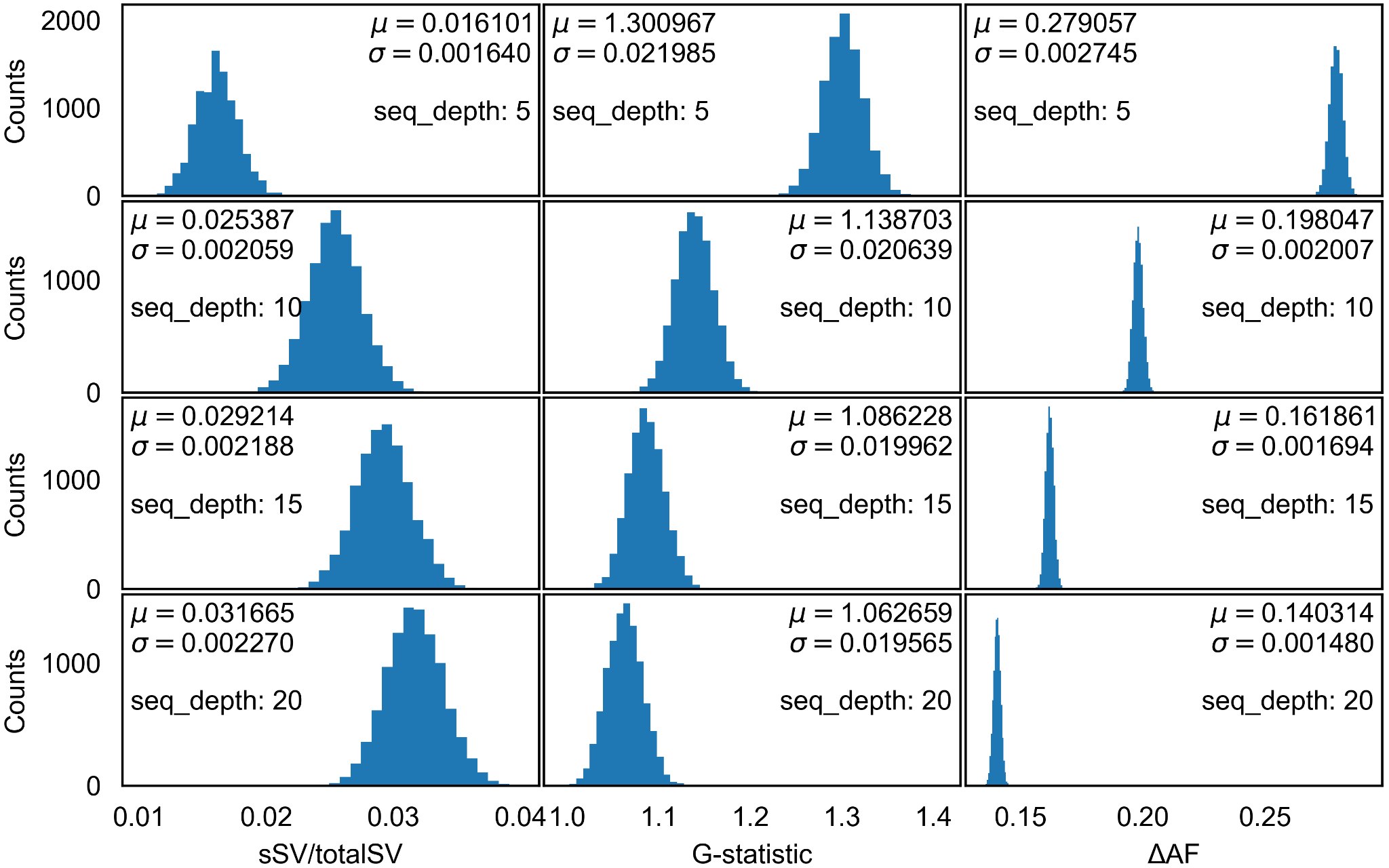

**Figure S2. Distribution of 10000 values of sSV/totalSV, *G*-statistic, and ΔAF obtained via simulation under the null hypothesis** (all SVs are not associated with the trait). Conditions: the average sequencing depths are 5, 10, 15, and 20, from top to bottom; the coefficient of variation (sequencing depth) is 0.4; the number of SVs per sliding window is 6000.
